# Single-Cell Atlas of CD4+ T-Helper 2 Cells Reveals Distinct Cytokine Functions Across Diseases

**DOI:** 10.1101/2025.09.20.677532

**Authors:** Son Vo, Maria Suprun, Han Do, My Ngo, Rajesh Rao, Saurabh Vishnu Laddha, Marta E Polak, Calixte Monast, Son Pham

**Affiliations:** BioTuring, 4445 Eastgate Mall, San Diego, California 92121, USA; Johnson & Johnson, 1400 McKean Road, Spring House, PA, 19477, USA

## Abstract

Despite the prevalence of autoimmune diseases, treatment options for specific conditions remain limited. These diseases are driven by three types of inflammatory pathways, each orchestrated by different T-helper cell subsets; however, a comprehensive atlas capturing their diverse functions across conditions and tissues is lacking. One of these subsets, CD4^+^ T-helper type 2 (Th2) cells, is a relatively infrequent immune cell population that plays a central role in orchestrating type 2 inflammation. In this study, we developed the first comprehensive CD4^+^Th2 single-cell atlas by analyzing the BioTuring database, which has data from over 300 million cells. This atlas integrates data from 52 studies, comprising a total of 39,243 cells derived from 9 major tissues, representing over 30 distinct diseases. Leveraging this atlas, we confirmed a rarely recognized CD4^+^Th2 cell population co-expressing *IL22* and *IL13* observed exclusively in atopic dermatitis and not in other allergic conditions. By integrating single-cell data with other “omics” technologies, our analysis suggests distinct downstream effects of *IL22* and *IL13*, highlighting the potential for combination therapy to enhance treatment outcomes for AD patients. Additionally, by utilizing data derived from the wide range of diseases, tissues, and time points included in this atlas, we re-confirmed the unique connection between *IL9* expression and conditions related to allergen exposure. Pseudotime analysis detailed CD4^+^Th2 cytokine dynamics upon allergen stimulation, showing a shift from early *IL2* and *IL4* expression to later *IL9* and *IL5*, with *IL13* consistently expressed, and further revealed that *IL9* production was transient. Collectively, these findings reveal distinct, context-dependent CD4^+^Th2 cytokine functions, opening the potential for identifying novel therapeutic strategies for allergic and inflammatory diseases.

## 1. Introduction

CD4^+^ T-helper type 2 (Th2) cells play a central role in type 2 immune responses, primarily targeting extracellular parasites. Current understanding of CD4^+^Th2 biology largely revolves around well-characterized cytokines such as IL4, IL5, and IL13, which are crucial for recruiting eosinophils and basophils to sites of infection^1^. However, overactivation of CD4^+^ Th2 cells has been implicated in chronic allergic inflammation, contributing to prevalent diseases like atopic dermatitis and asthma^2,3^. While existing therapies targeting these cytokines, such as dupilumab, have shown efficacy, not all patients achieve remission, suggesting involvement of additional, yet uncharacterized, factors/pathways in Th2-related diseases ^4^.

Despite their importance, the heterogeneity and functional diversity of CD4^+^Th2 cells across tissues and disease conditions remain poorly understood. This challenge arises partly from the limited availability of markers that can reliably differentiate Th2 cells from other T-helper cell subsets, relying mostly on cytokine secretion^5^. In addition, the low frequency of CD4+Th2 cells in peripheral tissues further complicates their detection and analysis. Moreover, the plasticity of different T-helper subsets, which allows for transition between different classes, such as T follicular helper to CD4+Th2^6^ or regulatory T cells to Th17^7^, poses additional challenges for identifying classes when a T cell expresses more than one type of marker.

To address this gap, we constructed the first integrated CD4+Th2 single-cell atlas, encompassing diverse tissues and a wide range of conditions to systematically study CD4+Th2 heterogeneity. Through detailed analysis, we uncovered distinct tissue- and condition-specific properties of CD4+Th2 cells, identifying *IL22* and *IL9* as disease-specific factors involved in atopic dermatitis and allergen-driven conditions, respectively.

## 2. Results

### 2.1. An integrated CD4+ Th2 atlas

We constructed a CD4+Th2 atlas utilizing the BioTuring single-cell database, which has data for over 300M cells (Fig. 1a). First, to identify consensus marker genes for the CD4+Th2 population, we reviewed each study that characterized CD4+Th2 cells, following annotations provided by the authors. We compiled a comprehensive list of the marker genes used in these studies to differentiate CD4+Th2 cells from other subtypes, and classified a gene as a consensus CD4+Th2 marker if it was consistently used for Th2 annotation in more than three studies. To ensure specificity, we excluded marker genes that were used to annotate other cell types in more than three studies. The final CD4+Th2 marker panel included *IL4, IL5, IL9, IL13, IL17RB, PTGDR2, HPGDS*, and *IL1RL1*. A CD4+ T cell was classified as a Th2 subset cell if it expressed at least one of these markers and highly expressed GATA3. Using this panel, combined with markers from other cell types as the negative panel, we applied a cell-type prediction algorithm to identify CD4+Th2 cells in pre-existing individual datasets, which were then integrated to construct the atlas (see Methods for details).

**Figure 1.**
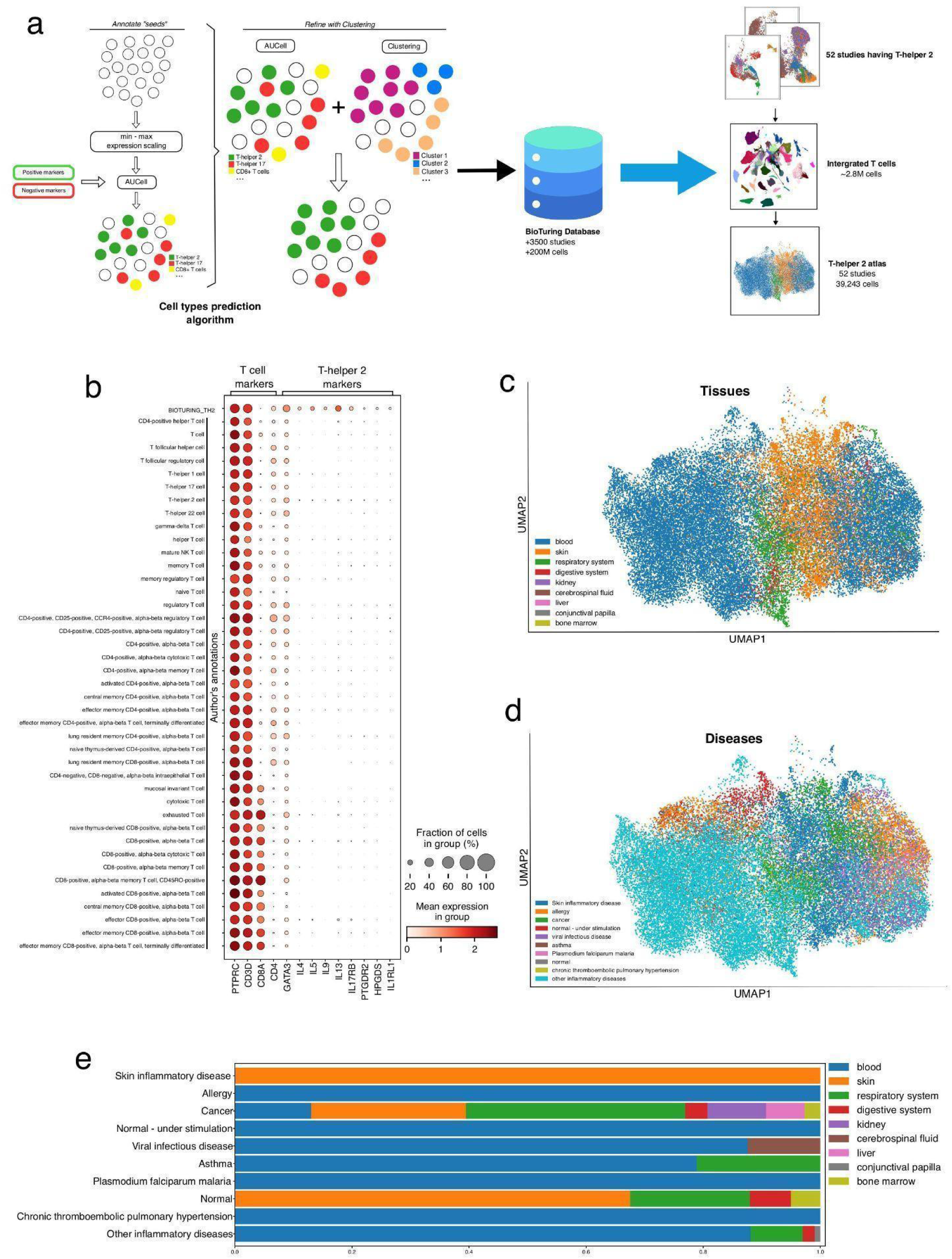
CD4+Th2 atlas overview. **a.** Summary of the cell type prediction algorithm and diagram of building the CD4+Th2 atlas from the BioTuring single-cell database. **b.** Dotplot visualizing expression of T cell marker genes and consensus Th2 marker genes (in columns) between BioTuring’s CD4+Th2 and the authors’ annotation (in rows) in the CD4+Th2 cell atlas. **c.** Distribution of major tissue types across CD4+Th2 cell atlas on UMAP. **d.** Distribution of conditions across CD4+Th2 cell atlas **e.** Proportions of tissues of origin in each condition

Applying these criteria to the BioTuring single-cell database, which is derived from over 3,500 studies and has data from more than 300M cells, we identified 52 datasets with at least 50 CD4+Th2 cells represented in each (Supplementary Table 1). Over 2.8M T cells were extracted from selected datasets, however, only 39,243 were classified as CD4+Th2. We then visualized expression of our panel of CD4+Th2 marker genes on the integrated T-cell atlas, comparing the BioTuring-Th2-derived set with the authors’ annotations (Fig. 1b). All the T cells expressed *PTPRC* and *CD3D*, which are established T-cell markers. While BioTuring-Th2 set showed expression of CD4+Th2 marker genes (*IL4, IL5, IL9, IL13, IL17RB, PTGDR2, HPGDS*, and *IL1RL1*) in the identified Th2 cell subset, there was minimal expression of these genes in other

T-cell subsets, indicating the specificity of our annotated atlas and confirming the rarity of the Th2 population (Fig 1b). The accuracy of our CD4+Th2 classification on different sequencing technologies is shown in Fig. S1a. The final CD4+Th2 atlas contains a set of 39,243 cells, encompassing 9 different major tissues and 9 major conditions, as well as healthy controls (Fig. 1c, d).

As shown in Fig. 1c, e, the atlas includes CD4+Th2 cells from a diverse range of diseases (e.g., allergy, asthma, cancer, infectious diseases, and skin inflammation) and healthy controls, as well as tissues (e.g., blood, skin, respiratory system, digestive system, kidney, cerebrospinal fluid, liver, and bone marrow). These broad categories encompass immunologically heterogeneous conditions with varying degrees of Th2 involvement. For example, among skin inflammatory diseases, Th2 cytokines (e.g., IL13) are prominently expressed in atopic dermatitis and prurigo nodularis, but not in Th17-driven conditions such as psoriasis^8^. Similarly, expression of Th2 markers varies across asthma subtypes and tumor types. To ensure immunological relevance, our downstream analyses focus on representative Th2-enriched contexts within each disease category. For clarity and consistency, we retain the use of broader category terms (e.g., asthma, cancer, skin inflammation) throughout the manuscript, with the understanding that they refer specifically to Th2-associated subsets, unless otherwise noted.

Some conditions had limited sampling sites; for example, all allergy samples were derived from blood, and all skin inflammatory samples came from skin tissue. Additionally, the atlas includes CD4+Th2 cells activated in vitro using established T-cell activation methods (PMID35618845). Beyond inflammatory and autoimmune diseases, the representation of CD4+Th2 cells derived from cancer was limited. Among the 11,330,316 T cells profiled from cancer samples in the BioTuring’s single-cell database^9^, only 2,761 (∼0.024%) were classified as CD4+Th2 cells, indicating a limited role of this T-helper subset in cancer. Of the 39,243 CD4+Th2 cells identified, only 682 (1.71%) originated from healthy samples, primarily from the skin and respiratory system, highlighting their rarity in non-disease conditions.

Following batch correction for study and assay technology, CD4+Th2 cells within the atlas tended to cluster based on their tissue of origin (Fig. 1c). CD4+Th2 cells from solid tissues clustered separately, with skin-derived CD4+Th2 cells being dominant and forming distinct clusters, suggesting potential tissue-specific properties of CD4+Th2 subsets.

### 2.2. Th2 Heterogeneity across Different Conditions

To explore Th2 heterogeneity across different diseases, we first visualized the expression of CD4+Th2 marker genes in major disease classes (Figure 2A). In allergy (peanut allergy, non-asthmatic subjects allergic to house dust mites) and asthma (allergic asthmatic patients, asthmatic subjects allergic to house dust mites), CD4+Th2 cells showed elevated expression of *IL4, IL5, IL9,* and *IL13*, highlighting their role in immune cell recruitment and inflammation^10–12^. In contrast, CD4+Th2 cells in Th2-associated inflammatory skin diseases predominantly expressed IL13, with minimal expression of the other three cytokines. We then visualized expression of the CD4+Th2 panel of markers across multiple studies, separated by tissue type. We observed that while *IL4, IL5,* and *IL13* showed expression in both blood and solid tissues, *IL9* was predominantly expressed in blood, with only minimal presence in solid tissues (Fig. S1b). In cancer, the expression levels of *IL4, IL5,* and *IL13* were low, while IL9 was undetectable. Upon investigation into more detailed conditions, we found that eosinophilic esophagitis (EoE), food allergy (FA), asthma/allergic respiratory diseases (A/ARD), and AD were the most extensively represented conditions in the atlas, based on cell numbers, samples, and datasets (Fig. 2b). As these were common inflammatory atopic diseases, these findings emphasize the central role of CD4+Th2 cells in inflammatory and allergic responses.

**Figure 2.**
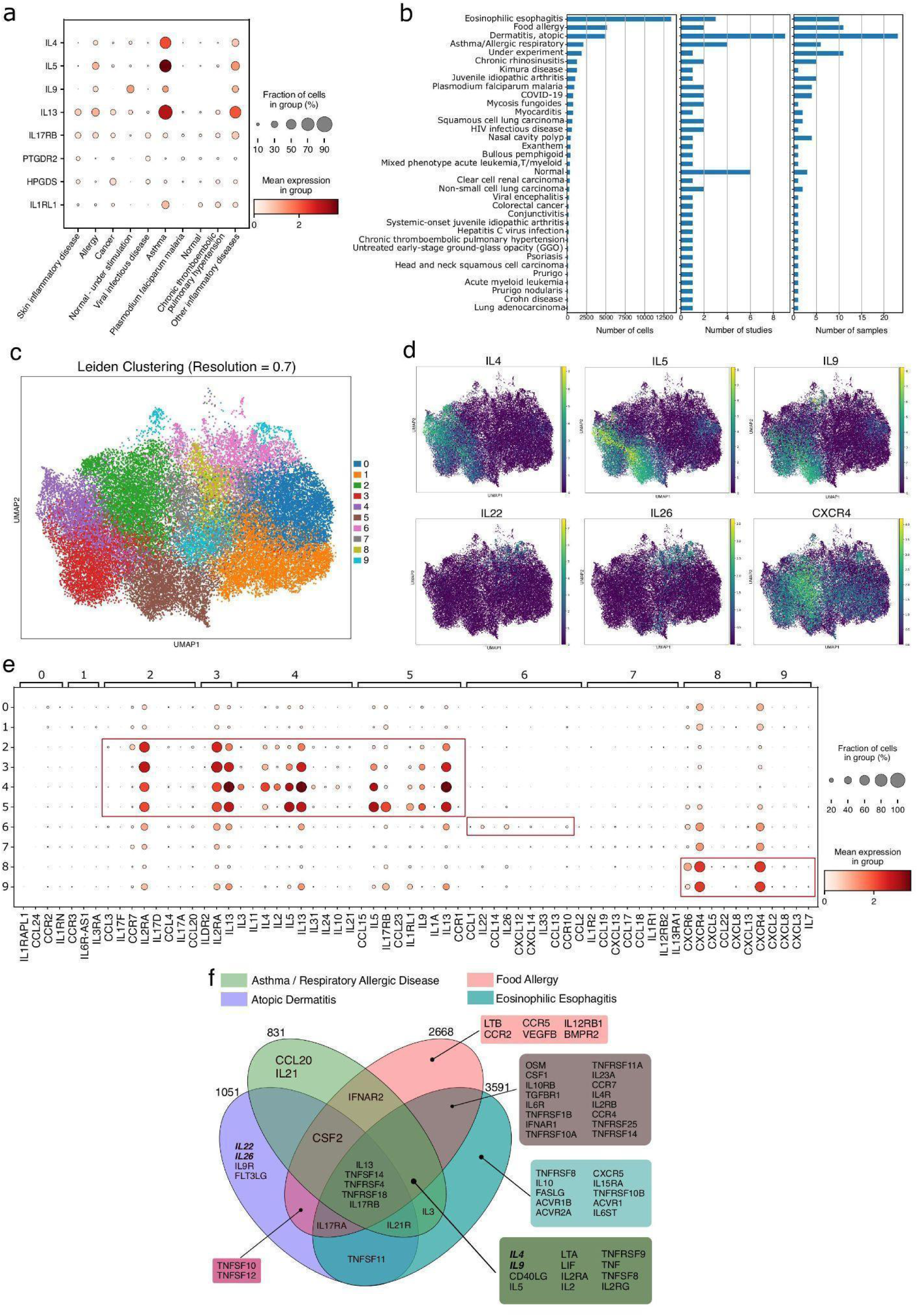
CD4+Th2 gene expression profiles across different conditions. **a.** Expression of CD4+Th2 cell markers between different groups of conditions, with enrichment of interleukin genes in asthma, allergy, and other inflammatory diseases. **b.** Number of cells, studies, and samples included for each condition in the CD4+Th2 cell atlas. **c.** Unsupervised Leiden clustering with resolution=0.7 on scVI embedding. **d** Expression of *IL4, IL5, IL9, IL22, IL26*, and *CXCR4* were visualized on UMAP. **e.** Dotplot showing the expression and coverage of the top differentially expressed cytokines and chemokines of each cluster. **f.** Venn diagram showing the overlap of differentially expressed genes in different Th2-associated diseases compared to healthy controls. Genes shown in the diagram are from the “KEGG CYTOKINE-RECEPTOR INTERACTION” pathway.

We then applied unsupervised clustering to identify sub-populations within the CD4+Th2 atlas (Fig. 2c), followed by differential gene expression analysis to extract top genes associated with each cluster. To focus on the immune regulatory roles of CD4+Th2, we narrowed the gene list (adjusted p-value < 0.05) to cytokines and chemokines. This approach revealed three major sub-populations (Fig. 2e). The first group (clusters 2, 3, 4, and 5) showed strong expression of *IL2RA, IL4, IL5, IL9*, and *IL13*. In contrast, cluster 6 represented a distinct group marked by elevated *IL22, IL26*, and *CCR10* expression, with minimal expression of cytokines from group 1. The third group (clusters 8 and 9) was characterized by a dominant chemokine profile, including *CXCR4 and CXCR6*. In terms of disease distribution, the first group was enriched in Kimura disease, EoE, FA, and A/ARD, whereas the second group was predominantly associated with AD.

To further understand the diversity in CD4+Th2 cell-driven inflammatory allergic mechanisms, we performed differential expression analysis comparing AD (n=9 datasets), A/ARD (n=4 datasets), FA (n=2 datasets), and EE (n=3 datasets) against normal control datasets (n=6). Given that CD4+Th2 cells mediate their functions through cytokines and their receptors, we focused on identifying cytokine- and receptor-related genes within the “KEGG CYTOKINE-RECEPTOR INTERACTION” pathway that are differentially expressed in these four conditions (Fig. 2f). Consistent with findings across major disease classes, *IL13* was elevated in CD4+Th2 cells in all conditions compared to controls. In contrast, *IL4, IL5*, and *IL9*, although previously defined as potential markers for CD4+Th2 cells, were predominantly expressed in FA, EoE, and A/ARD. Interestingly, AD did not exhibit high levels of expression of well-known Th2-associated cytokines such as *IL4* or *IL5,* but uniquely expressed *IL22* and *IL26*.

Due to the limited set of different tissue samples across these four diseases available in the atlas – predominantly skin in AD and blood in A/ARD, FA, and EE (Fig. 3a), we considered the possibility that tissue type, rather than disease state, could be the primary driver of cytokine expression, in some cases. To investigate this, we examined *IL4, IL5*, and *IL9* expression across all diseases sampled from blood (Fig. 3b) and *IL22* and *IL26* expression across all diseases sampled from the skin (Fig. 3c) in the Th2 atlas. Apart from A/ARD, FA, and EE, only Kimura disease exhibited elevated *IL4, IL5,* and *IL9* expression, suggesting that not all the CD4+ Th2 cells in blood express these cytokines. Similarly, in addition to AD, expression of *IL22* and *IL26* was observed only in mycosis fungoides and prurigo nodularis. While prurigo nodularis also showed strong expression of these cytokines, the sample contained fewer than 100 cells, indicating the signal may stem from background expression. Collectively, these observations suggest that the elevated expression of *IL4, IL5, IL9, IL22*, and *IL26* is driven by disease type rather than the tissue of origin.

**Figure 3.**
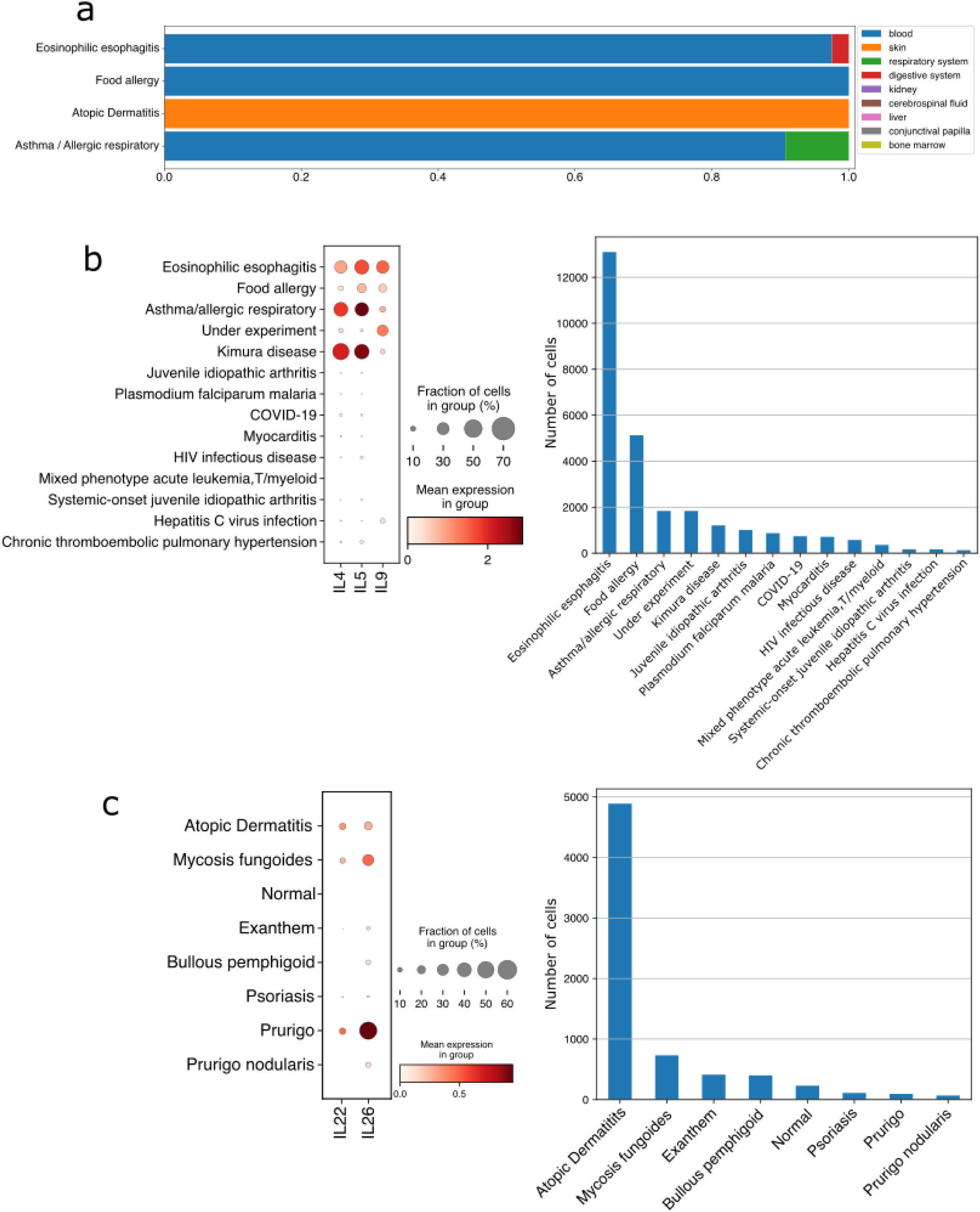
Cytokine expression in CD4+ Th2 cell atlas is disease-driven rather than tissue-dependent. **a.** Distribution of tissues in EE, FA, AD, and A/ARD. **b.** Expression levels of *IL4*, *IL5,* and *IL9* in all diseases sampling from blood in the Th2 cell atlas. Elevated expression was only observed in A/ARD, FA, EE, and Kimura disease, suggesting these cytokines are not associated with blood-derived Th2 cells. **c.** Expression of *IL22* and *IL26* in all diseases sampling from skin in the Th2 cell atlas. Beyond AD, notable expression was observed only in mycosis fungoides and prurigo nodularis, but with low cell counts, making it difficult to distinguish from background signals. This observation supports a disease-specific, rather than tissue-specific, pattern of *IL22* and *IL26* in CD4+Th2 cells.

### 2.3. Th2 cells co-express IL22 and IL13 in AD

IL22 was uniquely upregulated in AD compared to healthy controls (Fig. 2f and 3c), with prominent co-expression of IL13 (Fig. 4a–c). To quantify this, we analyzed the number of cells expressing *IL13, IL22*, or both across all conditions (Fig. 4d). A cell was classified as expressing a gene if at least one raw count was detected in its expression profile. AD displayed a high percentage (23.81%, n=424) of cells co-expressing *IL13* and *IL22*, confirming this as a unique feature of AD. This pattern was consistently replicated in four different datasets in the atlas (Fig. 4e).

**Figure 4.**
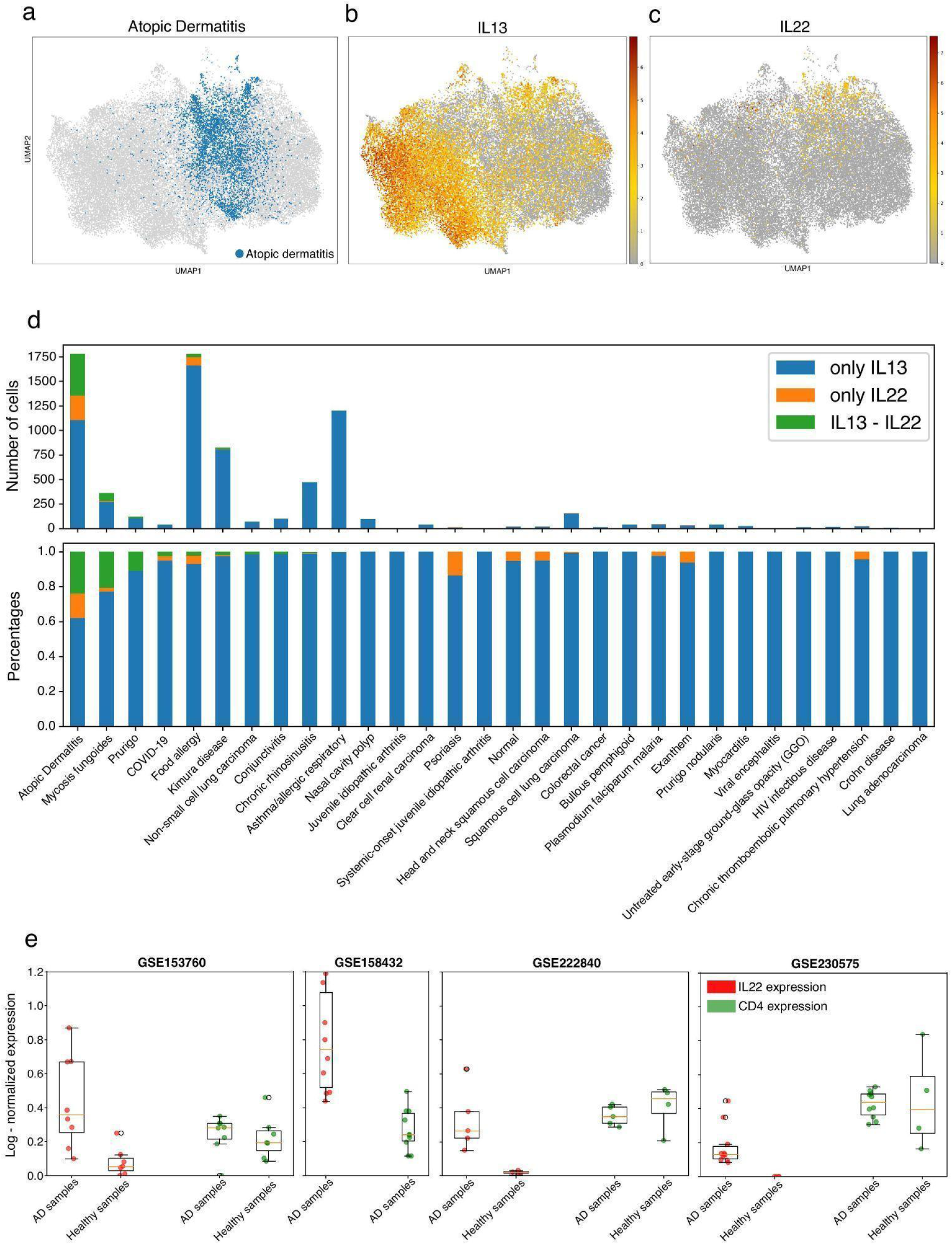
IL22 and IL13 Coexpression in Atopic Dermatitis. **a.** Distribution of cells from atopic dermatitis disease, along with *IL13* **(b)** and *IL22* **(c)** expression embedded on the same UMAP. Co-expression of *IL13* and *IL22* was primarily observed in atopic dermatitis. **d.** Number (top) and percentage (bottom) of cells expressing *IL13* and *IL22* across conditions. Atopic Dermatitis showed the largest number and ratio of co-expressed cells. **e.** Mean expression (log-normalized values) of *IL22* (red) and *CD4* (green) in CD4+ T cells in four studies that included atopic dermatitis ; each dot represents an individual batch/sample. All studies show a consistently elevated level of *IL22* expression in AD compared to healthy controls, while *CD4* expression, as a control, remained similar between AD and healthy controls.

We next investigated the role of *IL22* in AD by comparing its expression in other types of inflammatory skin diseases. We observed that *IL22* was co-expressed with *IL17A* (a Th17 marker) in psoriasis (Fig. 5a). Despite their distinct immune response types and T-helper cell origins, both AD and psoriasis exhibited elevated *IL22* expression. Since both diseases share histological and clinical features such as epidermal hyperplasia and skin inflammation, we hypothesized that *IL22* contributes to features of skin inflammation symptoms in AD independently of *IL13*.

**Figure 5:**
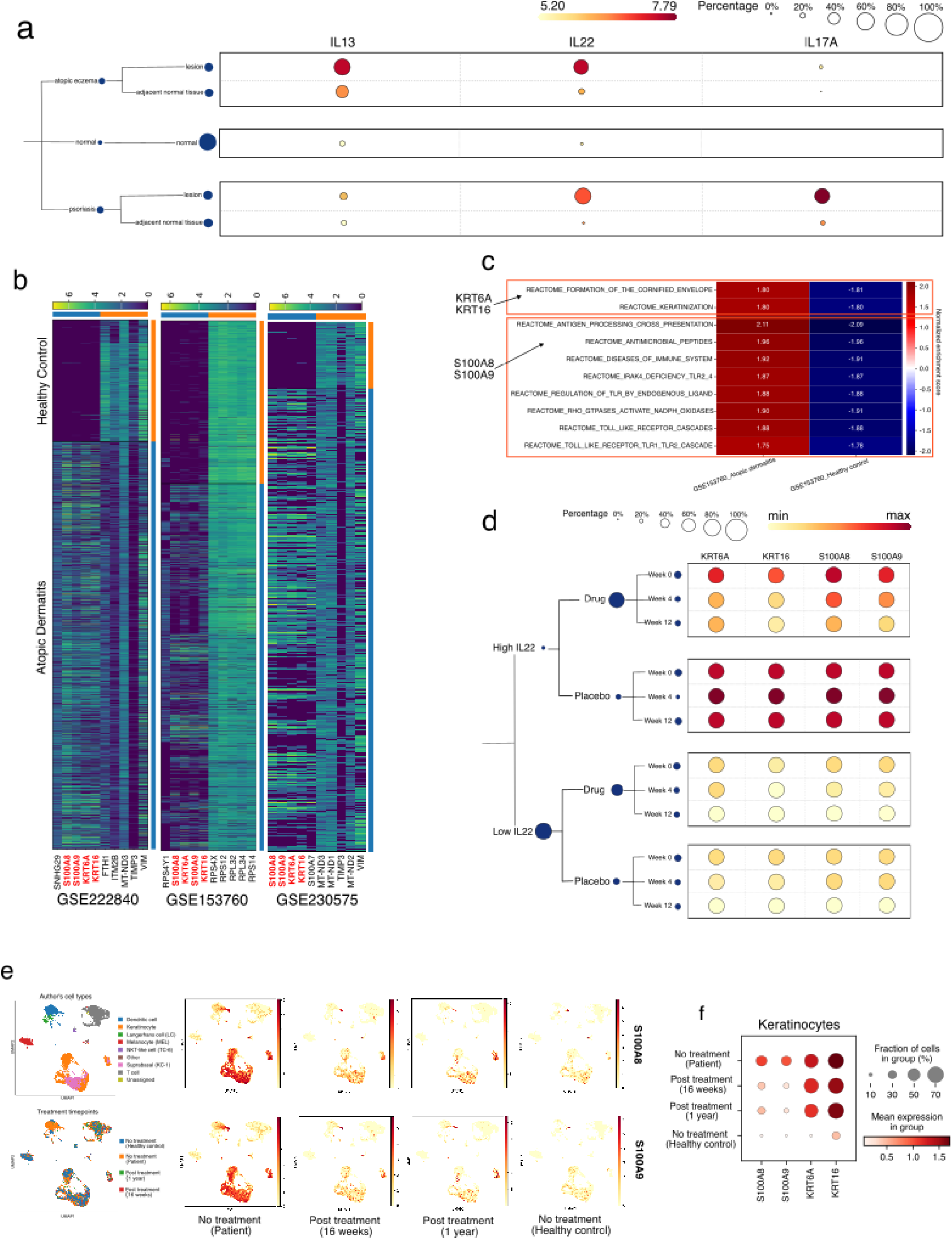
Downstream function of IL22 and IL13 in Atopic Dermatitis. **a.** Expression of *IL13, IL22,* and *IL17A* in atopic dermatitis, psoriasis, and healthy controls. *IL22* was still expressed when lacking either *IL13* (in psoriasis) or *IL17A* (in AD), proving its independent roles. **b.** Differential gene expression analyses between AD patients and healthy controls; all three studies showed the same top-upregulated genes in AD: *KRT6A, KRT16, S100A8*, and *S100A9*. **c.** GSEA heatmap showing the upregulated keratinization pathway related to *KRT6A, KRT16, S100A8,* and *S100A9* in AD patients compared to healthy controls. **d.** Fezakinumab reduces levels of *KRT* and *S100A* gene expression in atopic dermatitis patients with high IL22 baseline. **e, f.** Dupilumab reduces S100A gene expression and has a minor effect on KRT genes in keratinocytes from atopic dermatitis patients.

We first performed differential gene expression (DEG) analysis comparing AD and normal skin samples. To avoid technical variation between datasets, we conducted DEG analyses separately for each dataset. In three datasets (GSE153760, GSE222840, and GSE230575), the top upregulated genes in AD were *KRT6A, KRT16, S100A8,* and *S100A9* (adjusted p-value < 0.05) (Fig. 5b). Visualization of these genes revealed consistent upregulation across all cell types in AD compared to normal samples, with the most pronounced differences observed in keratinocytes (Fig. S2). To investigate the functional relevance of upregulation of these genes, we performed pathway enrichment analysis. AD samples exhibited upregulation of skin disorder pathways, including “Formation of the Cornified Envelope” and “Keratinization”, both of which include *KRT6A* and *KRT16* genes. In contrast, *S100A8* and *S100A9* genes are associated with pathways related to “Antimicrobial Peptides” (Fig. 5c). While antimicrobial peptides are pertinent to the innate immune system, their excessive upregulation could contribute to skin inflammation^13^. Bonafice et al. 2005 showed by RT-PCR that *S100A7, S100A8,* and *S100A9* mRNA levels increased in a dose-dependent manner in response to *IL22*; western blot analysis confirmed that IL22 stimulation led to elevated levels of S100A proteins^14^. We then set out to evaluate the relationship between *IL22* and *KRT6A, KRT16, S100A8*, and *S100A9* to understand the connection between *IL22* and AD endotypes.

IL22 inhibition by fezakinumab, an IL22-blocking biologic agent, reduced immune activation in the skin, as evidenced by decreased expression of *S100* genes, especially in the IL22-high expression population (GSE99802^15^). However, this study did not address the effects of IL22 inhibition on *KRT6A* and *KRT16*. We then reanalyzed this dataset by comparing the expression patterns of these genes across patients treated with fezakinumab versus placebo (Fig. 5d).

Consistent with findings on *S100* genes from the original publication, stratification by baseline IL22 levels was critical for detecting meaningful transcriptional changes in *KRT* genes following treatment. In AD patients with high baseline IL22 expression, fezakinumab effectively reduced levels of *KRT6A* and *KRT16* expression after 4 weeks of treatment. However, no significant effect was observed in AD patients with low IL22 baseline levels or those receiving a placebo.

The relationship between *IL13* and four upregulated downstream genes (*KRT6A, KRT16, S100A8*, and *S100A9)* was mentioned in Bangert et al. 2021^16^. Treatment with dupilumab led to a significant reduction in the coverage and expression levels of *S100A8* and *S100A9* in keratinocytes (Fig. 5e). However, there were only minor changes in *KRT6A* and *KRT16* expressions. These results indicate that dupilumab effectively mitigates the upregulation of antimicrobial peptide gene expression, but does not impact expression of hyperplasia-related keratin genes.

Leveraging the atlas helped identify the unique pattern of *IL13* and *IL22* co-expression in AD. However, our analyses proposed distinct downstream effects on AD pathology. Dupilumab, which blocks *IL4RA*, primarily influences expression of antimicrobial peptide genes with no significant effect on keratinization genes. In contrast, fezakinumab, an IL22 blocker, reduced expression of both keratinization and antimicrobial peptide genes. Based on these observations, we suggest a combination therapy targeting both cytokines could provide more comprehensive benefits in managing AD .

### 2.4. Temporal expression of IL9 in CD4+Th2 upon stimulation

In contrast to *IL22* being predominantly expressed in AD, *IL9* was expressed only in FA, EE, and A/ARD (Fig. 2f). To determine whether this was due to the difference in tissue type, we explored *IL9* expression in blood and bronchial tube samples across eight conditions, spanning allergen-induced to non-allergen-induced diseases (Fig. 6a). Both blood and bronchial tube samples from A/ARD exhibited high IL9 expression. In fact, all allergic conditions subjected to either natural or artificial stimulation expressed *IL9*, while non-allergic blood samples showed no *IL9* expression (adjusted p-value <0. 05). This suggests that production of *IL9* in CD4+Th2 cells might be a consequence of allergen exposure.

**Figure 6.**
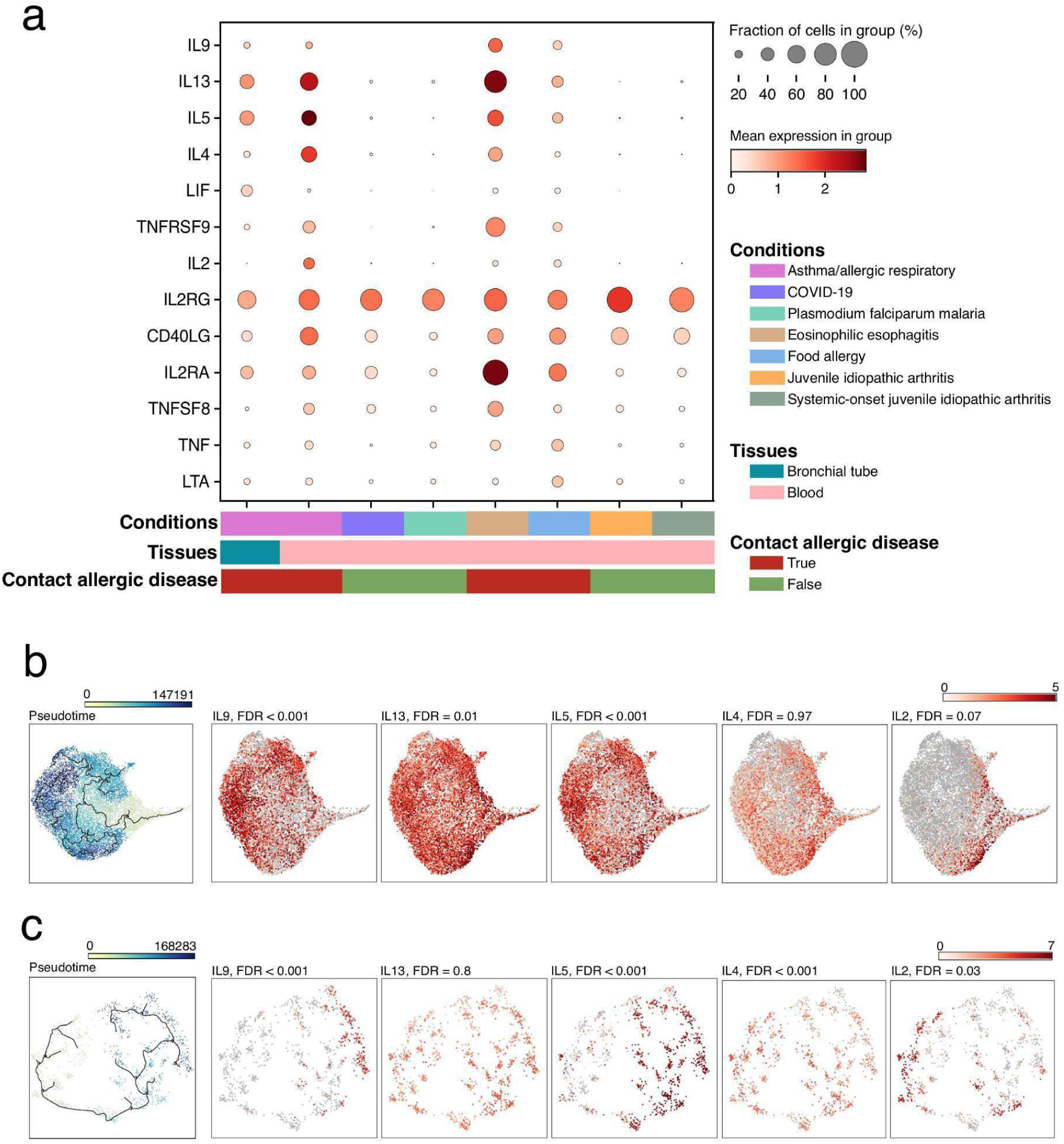
IL9+ CD4+Th2 Subpopulation Characterization. **a.** Dotplot showing upregulation of genes across different conditions in bronchial tube and blood samples. **b, c.** Pseudotime trajectory analysis and expression patterns of *IL9, IL13, IL5, IL4*, and *IL2* in **(b)** eosinophilic esophagitis (EE) (GSE175930, n = 13,100 Th2 cells) and **(c)** A/ARD (GSE146170, n = 1,176 Th2 cells). In both diseases, *IL13* maintained a consistent expression pattern throughout the trajectory, while *IL9* and *IL5* were enriched at one end and *IL4* and *IL2* at the opposite end.

To investigate *IL9* expression dynamics in Th2 cells, we performed pseudotime analysis on CD4+Th2 cells from contact allergic conditions in two datasets individually: blood samples from A/ARD patients stimulated with house dust mites for 6 hours (GSE146170, n=1,176 cells) and blood samples from EE patients stimulated with anti-CD3/CD28 for 6 hours (GSE175930, n=13,100 cells). Both analyses showed consistent interleukin expression patterns. Specifically, we found a transition from high *IL2* and *IL4* expression to high *IL9* and *IL5*, while *IL13* remained consistently expressed throughout the trajectory (Fig 6b, c). Given that *IL2* and *IL4* are typically associated with the early stages of CD4+Th2 polarization and activation, it is likely that *IL9* and *IL5* are expressed at later stages after stimulation.

To validate this hypothesis, we extended the analysis to include non-polarized CD4+ T cells, assuming pseudotime would progress from a naive to an activated CD4+Th2 state. Of the two datasets analyzed earlier, only the EE dataset (GSE175930) contained such cells. Therefore, we also included a bronchial tube asthma dataset (GSE193816) stimulated with house dust mite for 24 hours, despite the limited CD4+Th2 cell count (n = 373) in this dataset. Using non-polarized CD4+ T cells as the origin, both analyses revealed a temporal transition in CD4+Th2 cells from being *IL9* and *IL5* negative to becoming *IL9* and *IL5* positive, with consistent expression of *IL13* (Fig. 7a–d). This observation was consistent with the finding in the original study that *IL9* expression was elevated specifically in asthmatic airways upon allergen stimulation^17^. Notably, we observed early production of *IL2* in the EE and blood datasets (Fig. 7c, d). Additionally, the bronchial tube asthma dataset showed a unique trajectory, with Th2 cells reverting from being *IL9* positive back to *IL9* negative at the pseudotime endpoint (Fig. 7d). When focusing on allergic datasets undergoing stimulation at different time points, we found a similar pattern: *IL9* expression increased initially but decreased over time (Fig. S4).

**Figure 7.**
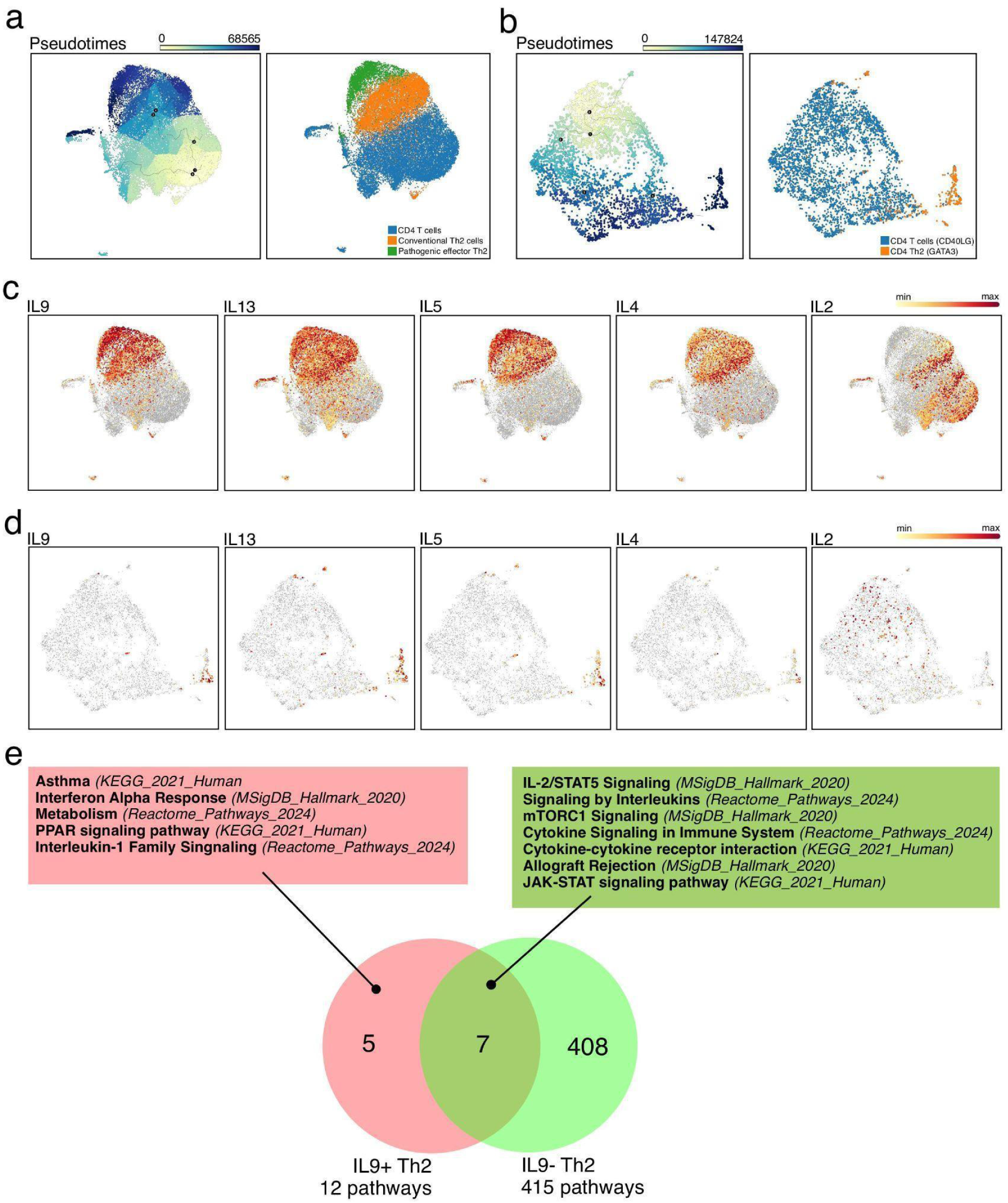
Pseudotime Trajectory and Pathway Enrichment Analysis of the IL9+ CD4+Th2 cell Subpopulation in EE and Asthma. **a-d.** Pseudotime trajectory analysis of CD4 T cells, including CD4+Th2 cells, from **(a)** EE (GSE175930, n = 30,635 cells) and **(b)** asthma (GSE193816, n = 4,503 cells) datasets, with corresponding expression of *IL9, IL13, IL5, IL4,* and *IL2* shown in **(c)** and **(d)**, respectively. **c, d.** Given that *IL2* and *IL4* are typically associated with the early stages of CD4+Th2 polarization and activation, early production of IL2 was observed in the EE blood datasets. **d.** Th2 cells reverting from IL9 positive back to IL9 negative expression at the pseudotime endpoint **e.** Pathway enrichment analysis comparing *IL9*-positive (IL9+ Th2) and *IL9*-negative (IL9- Th2) Th2 cells. The Venn diagram highlights the overlap of significantly enriched pathways, with 5 pathways unique to *IL9+* CD4+Th2 cells, 408 pathways specific to *IL9-* CD4+Th2 cells, and 7 pathways shared between the two populations. Key enriched pathways are annotated for each category.

To better understand the mechanisms driving *IL9* expression upon allergen exposure, a pathway analysis was conducted using highly expressed genes from *IL9*+ and *IL9−* Th2 cells, comprising 7,298 and 12,569 cells, respectively, across all A/ARD, EE, and FA datasets. The analysis identified five pathways uniquely and significantly enriched in *IL9+* Th2 cells, several of which are linked to inflammatory responses, reinforcing *IL9*’s established role in modulating immune responses in allergic diseases such as asthma (Fig. 7e). Intriguingly, *IL9*+ CD4+Th2 cells were also enriched for the PPAR signaling pathway, a critical regulator of systemic metabolism and energy homeostasis^18^.

## 3. Discussion

This study leveraged a large-scale single-cell database to construct the first comprehensive atlas of human CD4+Th2 cells across diverse tissues and diseases. Addressing the known challenges of identifying and characterizing this relatively rare T cell subset, our integrated approach successfully identified nearly 40,000 Th2 cells, defined by a consensus marker panel. The atlas confirmed the scarcity of CD4+Th2 cells in healthy tissues and cancer, reinforcing their primary association with inflammatory and allergic conditions, particularly AD, EE, allergy, and asthma. Initial analysis revealed that CD4+Th2 cells cluster significantly by their tissue of origin, suggesting that tissue-specific imprinting influences their function, although our subsequent analyses pinpointed specific disease states as major drivers of distinct functional cytokine profiles.

This CD4+Th2 atlas contributes to our understanding of the diversity and specificity of this rare CD4+ T-cell subset. Further, this atlas provides a systematic, high-resolution view that enables comparisons across datasets and conditions, revealing context-dependent cytokine expression patterns that might otherwise be overlooked. Despite its relatively small size, the atlas has capacity to capture distinct pathogenic mechanisms across different tissues and disease states. Unsupervised clustering over the CD4+Th2 atlas revealed multiple sub-populations, each marked by unique cytokine and chemokine signatures. The atlas introduced novel observations, such as the independent downstream functions of *IL13* and *IL22* in atopic dermatitis, as well as the transient expression of *IL9* as a consequence of stimulation in allergy and asthma. As such, the Th2 atlas provides a resource for refining our understanding of Th2-mediated immunity and identifying new therapeutic opportunities in Th2-associated diseases.

## 4. Limitation of Study

Although the creation of this CD4+Th2 atlas represents a pioneering effort in characterizing the complexities of this immune subset, certain limitations must be acknowledged. Derived consensus marker panels based on the majority of authors’ CD4+Th2 annotations may introduce biases. The coverage across all tissue-condition combinations is uneven, with some diseases primarily sampled from single tissues. Furthermore, the rarity of CD4+Th2 cells, particularly in cancer and healthy states, limits the depth of analysis in these contexts. Specifically, our CD4+Th2 marker panel seems skewed toward activated CD4+Th2 cells, as these markers are primarily associated with cytokine secretion, potentially overlooking markers for memory CD4+Th2 cells. Inclusion of datasets from additional studies may help address these limitations in the future versions of this atlas.

## 5. Methods

### 5.1. Marker Gene Panels Identification for CD4+Th2

To accurately identify CD4+Th2 cells, we constructed a positive marker gene pool through an extensive review of scientific literature and single-cell data. In addition, we utilized BioTuring Venice^25^ to extract Th2-specific markers from BioTuring database annotations. To ensure both consensus and specificity, we scored each marker gene based on the authors’ Th2 annotations in existing datasets:

- True positive score: A gene was scored as true positive if it effectively defined the CD4+Th2 cluster without significant expression in other CD4+ T-cell subtypes across all datasets.

- False positive score: A gene was scored as false positive if it effectively defined the CD4+Th2 cluster in one dataset but showed significant expression in other cell types in other datasets.

We iteratively assessed all datasets containing CD4+Th2 cells, retaining genes with true positive scores greater than three and false positive scores less than three.

### 5.2. Seed Detection using AUCell

An AUCell^26^ score was computed for every cell in the BioTuring single-cell database to detect CD4+Th2 cells (referred to as “seeds”). This score represents the enrichment of a given gene set in an individual cell. This scoring system was used instead of assessing expression of the exact gene panel to address the drop-out issue, which is common in single-cell sequencing technologies.

A combination to define each cell type includes:

*combination = [positive panel, negative panel]*.

The positive panel of CD4+Th2 cells is defined as:

*positive panel = [CD4+ T cell markers] + [Th2 markers]*

*CD4+ T cell markers* include three genes: PTPRC, CD3D, and CD4. To address the gene drop-out issue, we created permutations of these markers. “PTPRC CD4” set was excluded, since it is used to detect myeloid cells instead of CD4+ T cells. Resulting three panels included:

1. PTPRC, CD3D, CD4
2. PTPRC, CD3D
3. CD3D, CD4

Following the approach above, *Th2 markers were separated into 9 permutation sets.* Finally, combining *[CD4+ T cell markers] and [Th2 markers],* we created a total of 3 x 9 = 27 positive panels for CD4+Th2 cell identification (table below).

We included one negative panel of cell markers. This included marker genes derived from other major cell types, such as Epithelial cells, Endothelial cells, Myeloid leukocytes, B cells, gamma-delta T cells, and CD8+ T cells, naive T cells, Th1, Th17, T follicular helper, and regulatory T cells.

| Positive panel | Negative panel |
| --- | --- |
| PTPRC, CD3D, CD4, IL4 | CD34, EPCAM, PECAM1, CD14, CD79A, CD8A, CD8B, TRDC, TRGC1, TRGC2, CD27, FCER1G, GZMA, LEF1, IFNG, CCR6, IL17A, IL17F, CXCR5, FOXP3 |
| CD3D, CD4, IL4 |  |
| PTPRC, CD3D, IL4 |  |
| PTPRC, CD3D, CD4, IL5 |  |
| CD3D, CD4, IL5 |  |
| PTPRC, CD3D, IL5 |  |
| PTPRC, CD3D, CD4, IL9 |  |
| CD3D, CD4, IL9 |  |
| PTPRC, CD3D, IL9 |  |
| PTPRC, CD3D, CD4, GATA3-AS1 |  |
| CD3D, CD4, GATA3-AS1 |  |
| PTPRC, CD3D, GATA3-AS1 |  |
| PTPRC, CD3D, CD4, IL17RB |  |
| CD3D, CD4, IL17RB |  |
| PTPRC, CD3D, IL17RB |  |
| PTPRC, CD3D, CD4, PTGDR2 |  |
| CD3D, CD4, PTGDR2 |  |
| PTPRC, CD3D, PTGDR2 |  |
| PTPRC, CD3D, CD4, IL13 |  |
| CD3D, CD4, IL13 |  |
| PTPRC, CD3D, IL13 |  |
| PTPRC, CD3D, CD4, HPGDS |  |
| CD3D, CD4, HPGDS |  |
| PTPRC, CD3D, HPGDS |  |
| PTPRC, CD3D, CD4, IL1RL1 |  |
| CD3D, CD4, IL1RL1 |  |
| PTPRC, CD3D, IL1RL1 |  |

Min-max scaling was applied on all cell type markers for each cell before AUCell calculation to ensure uniform capture rates for each gene when determining representative cell types.

**Seeds detection algorithm:**

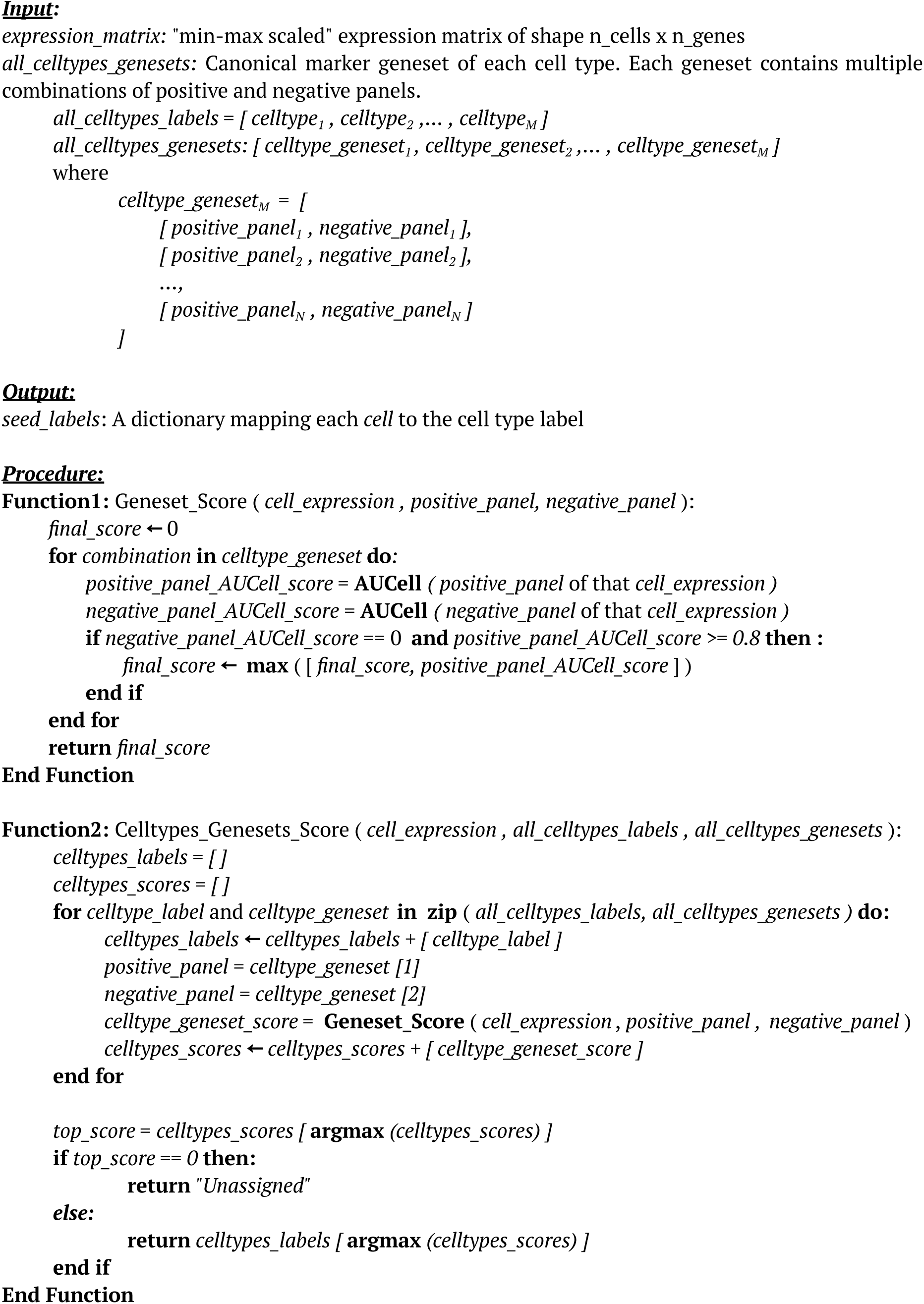
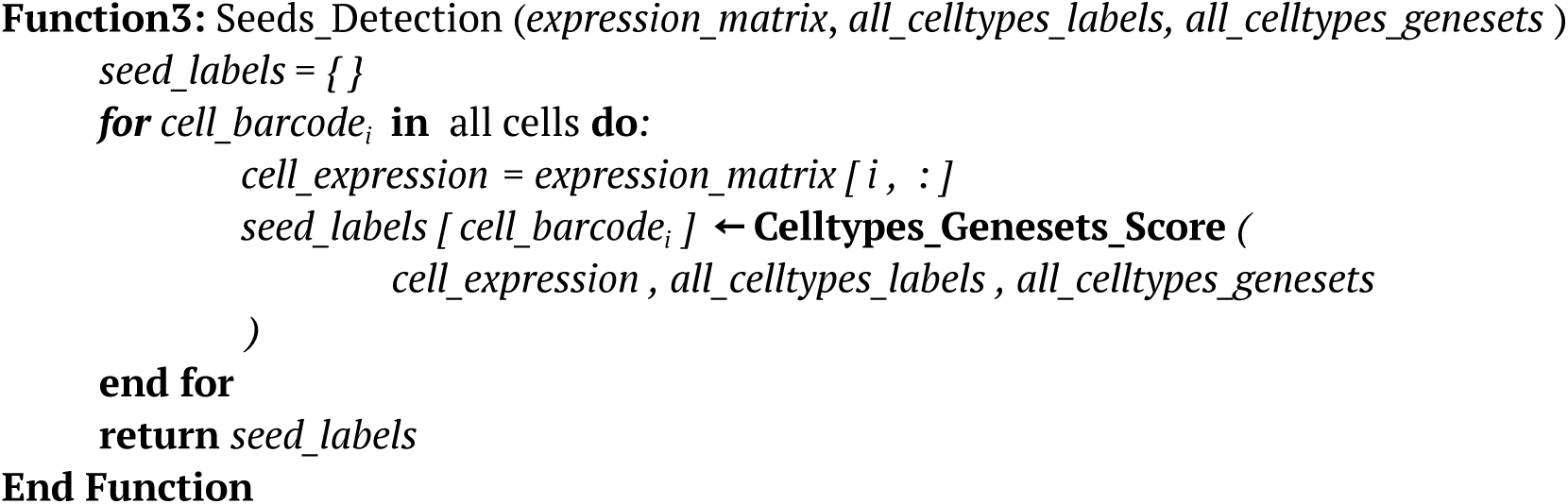

### 5.3. Seed Refinement using Iterative PCA, KNN, and Louvain Clustering

In this refinement process, we smoothed the identified seeds to their closed neighbor cells, which were determined by an integrated pipeline of iterative Principal Component Analysis (PCA)^28^, k-nearest neighbors (KNN)^29^, and Louvain clustering^30^. This step helped to mitigate the drop-out effects prevalent in single-cell datasets, while also discerning and removing false-positive seeds resulting from background noise expression.

We executed the pipeline independently within each sample across all studies, without employing batch correction methods. When performing PCA and Louvain clustering to annotate the cell types, clusters were correctly annotated only when all of its sub-clusters exhibited similar annotations. Any divergence from this pattern necessitated separation into smaller clusters to ensure accurate cell type annotations, the Louvain clustering algorithm was executed at high resolution to smooth the seeds effectively.

Given the focus on CD4+Th2 cells, all cell types apart from T cells resulting from seed detection were labeled as “Unassigned”. Throughout the smoothing process, clusters consisting solely of unassigned cells were iteratively eliminated, and PCA and the Louvain algorithm were rerun to improve separation between CD4+ T cell classes, thus improving Th2 annotation. A cluster was annotated as CD4+Th2 if its cell count ratio exceeded that of all other T cells. Finally, all the Th2 clusters from each of the different studies in the database were merged to create the final atlas.

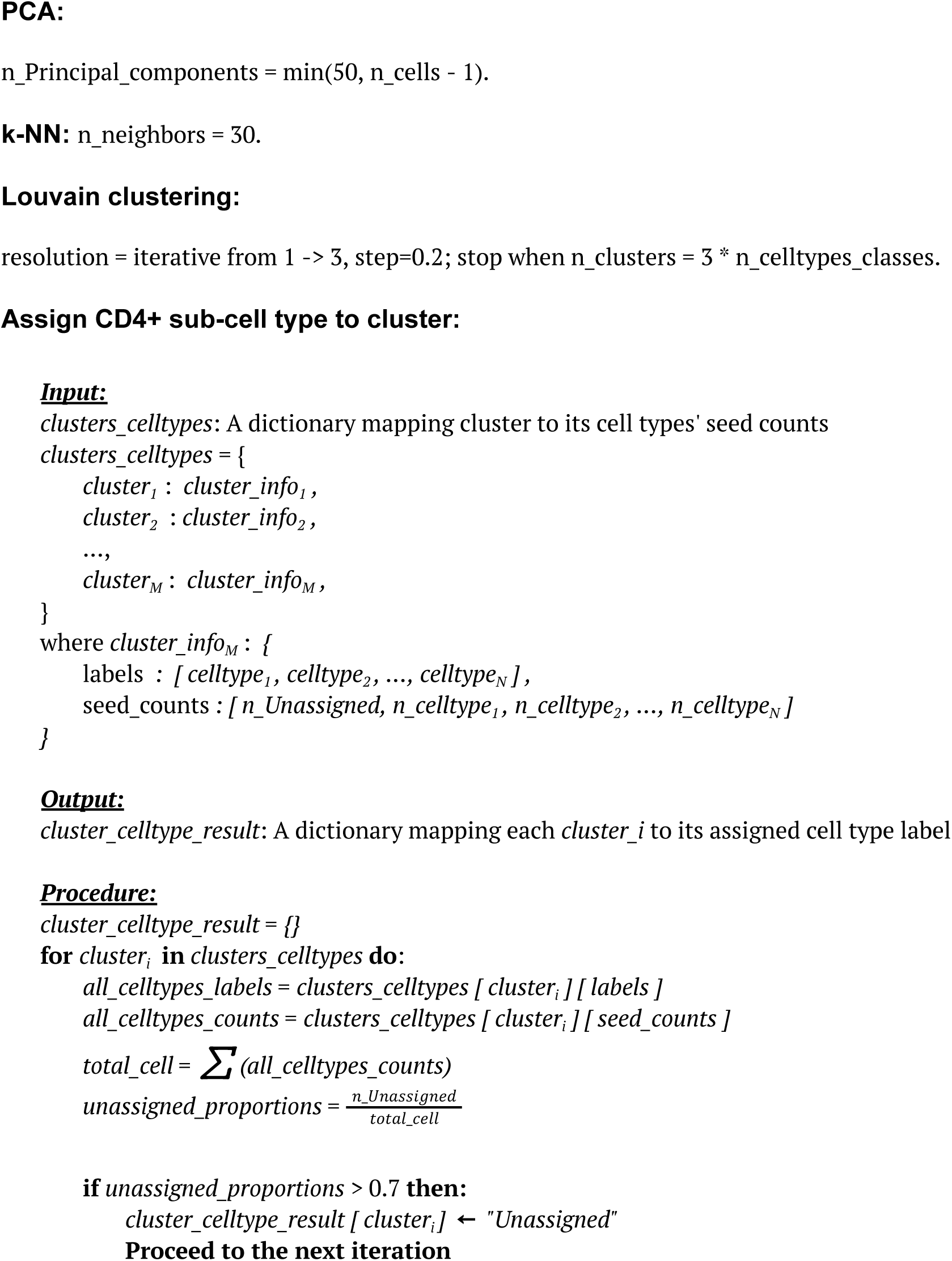
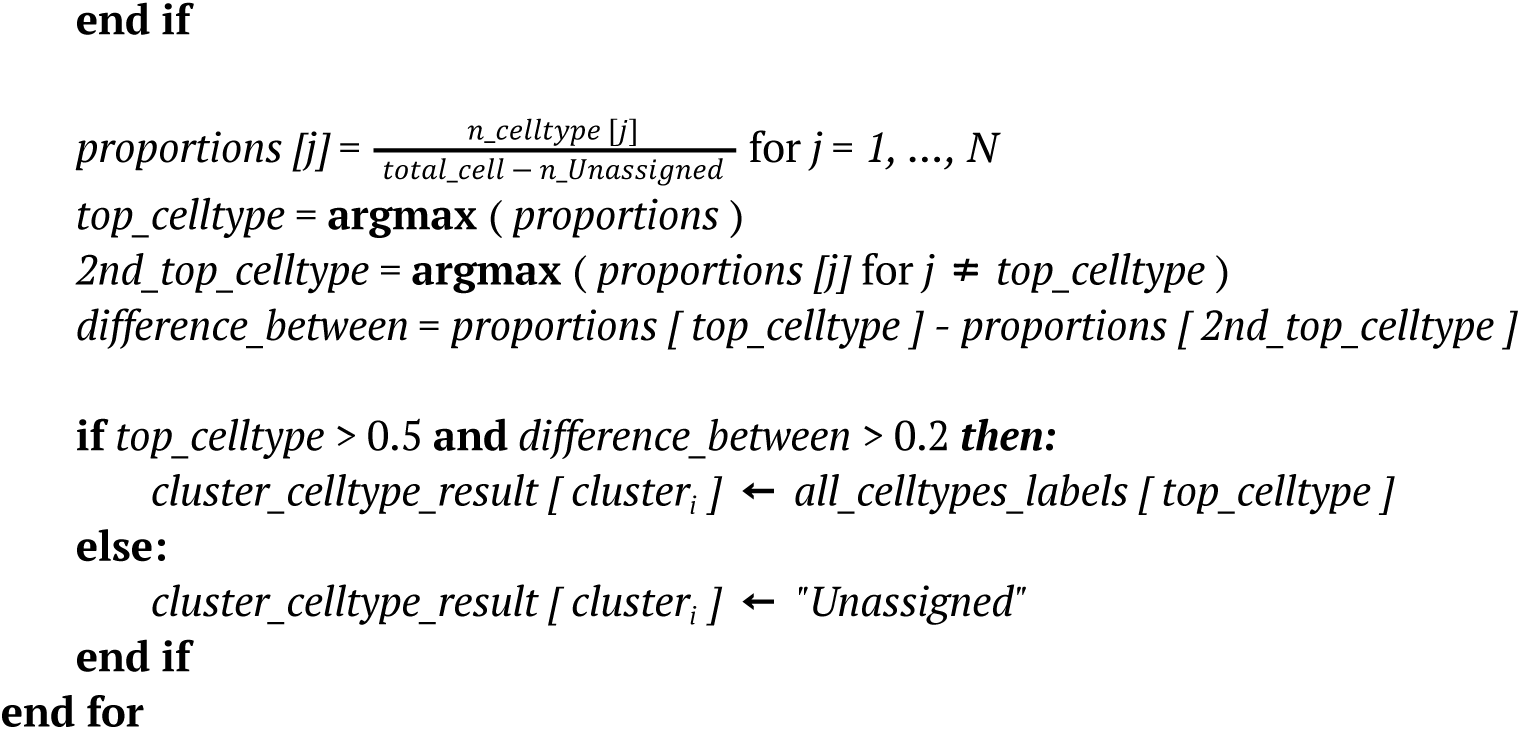

### 5.4. Preprocessing and Integrating for Atlas using scVI

Each selected dataset underwent individual preprocessing procedures strictly following the filtering parameters defined by the authors in the original papers (supplement table). All datasets included in the atlas were collected as raw integer counts, and the samples with fewer than 50 CD4+Th2 cells were excluded. The sequencing depth of all cells was then normalized by dividing the total counts over all genes of each cell by 10,000 and taking the natural logarithm of these values .

To ensure consistency of metadata terms across multiple datasets, the metadata of each dataset collected from each original publication, were carefully standardized based on EBI ontologies. For batch integration, we initially identified 1,000 highly variable genes (HVGs) using the seurat_v3 package implemented in Scanpy^31^. Subsequently, scVI^32^ was applied to the raw counts of this set of 1,000 HVGs. The integration process involved treating “Study ID” variable as the batch variable. The scVI method was run using specific parameter configurations: number of hidden dimensions: 128; number of layers: 1; number of latent dimensions: 30; gene likelihood: zero-inflated negative binomial (zinb; batch size: 128 and kl_weight: 0.1. The model was trained with 100 epochs to avoid over-batch correction, and all other parameters were set to default values. Finally, we computed a k-nearest neighbors graph with n_neighbor = 90 and applied UMAP^33^ using the scVI latent space for visualization.

### 5.5. Differential gene expression and Pathway analysis

Unless otherwise stated, Differential Gene Expression (DGE) analysis was performed using the BioTuring BBrowserX platform with the Venice statistical method. The analysis criteria included adjusted p-values < 0.05, |log2FC| > 0.5, Bonferroni correction for multiple comparisons, and a default min.pct threshold of 0.1 for gene coverage. The resulting gene scores were used as input for Decoupler^34^, while the differential gene list was put into GSEApy^35^ for pathway enrichment analysis. Enrichment tests incorporated pathways from the MsigDB^36^ (MSigDB_Hallmark_2020), KEGG^37^ (KEGG_2021_Human), GO^38,39^ (GO_Biological_Process_2023), and Reactome databases^40^ (Reactome_Pathways_2024). Pathways were considered significant if they had a threshold of adjusted p-values of < 0.05.

### 5.6. Trajectory analysis

Pseudotime trajectory analysis was performed using Monocle3 to investigate the temporal dynamics of interleukin expression in CD4+Th2 cells during allergen stimulation. To minimize batch effects from variations in condition, sequencing technology, and tissue, the analysis was performed on each dataset independently. Non-polarized CD4+ T cells were selected as the starting point to establish a naive-to-activated trajectory.

## Author contributions

SV created the pipeline, analyzed the data, prepared the figures, and wrote the manuscript. MN analyzed the data and wrote the manuscript. HD analyzed the data. MS, RR, SL, MP wrote the manuscript, designed the analytics. SP, CM conceived of the study, contributed to the creation of the algorithm, writing of the manuscript, and supervised the study.

## SUPPLEMENTARY DATA

**Supplementary Figure 1.**
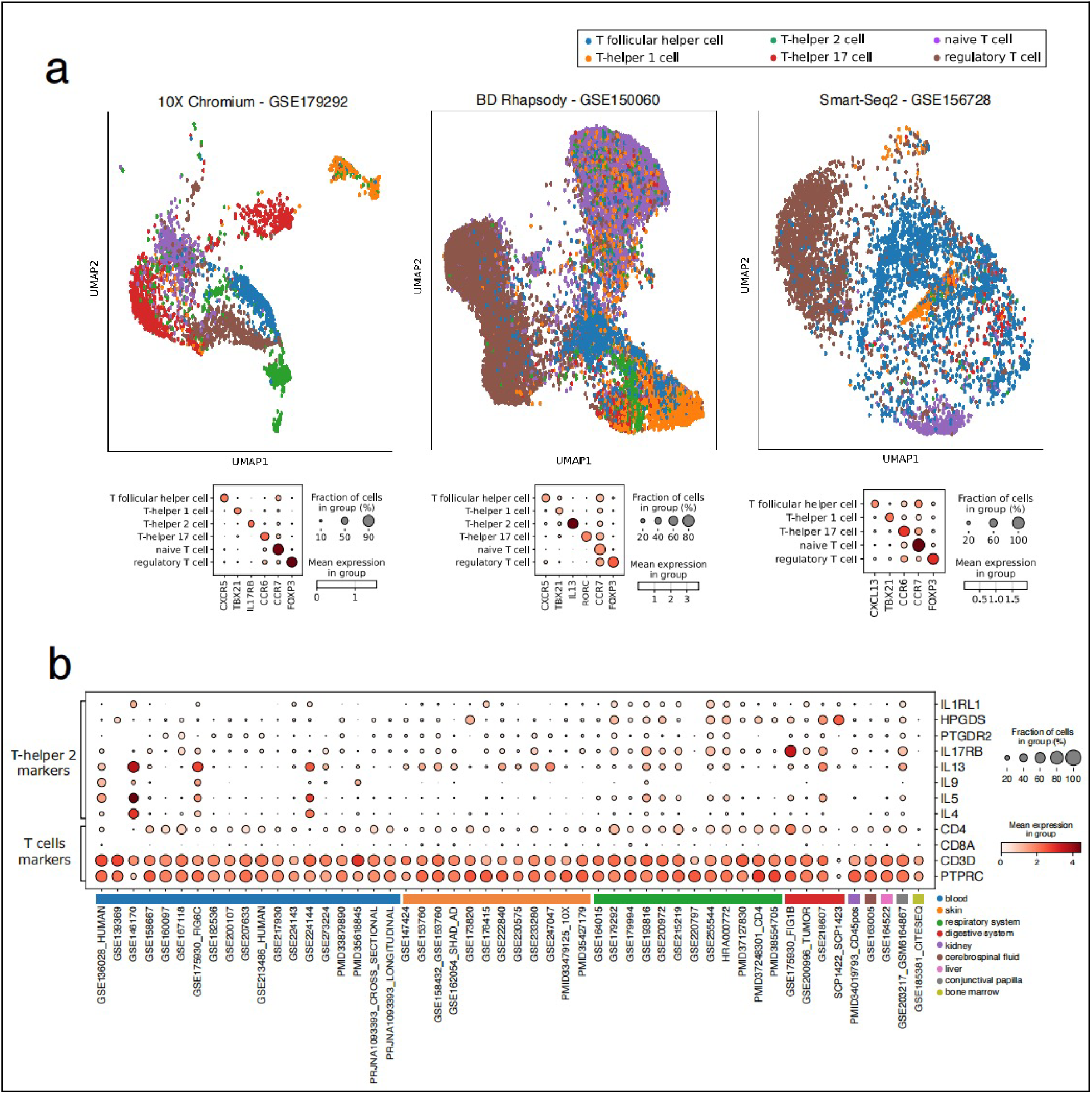
CD4+ Th2 cell type prediction results. **a.** CD4+Th2 cell type prediction results on different sequencing technologies. **b.** CD4+Th2 cell marker expression across individual studies included in the CD4+ Th2 cell atlas, grouping by tissue of origin.

**Supplementary Figure 2.**
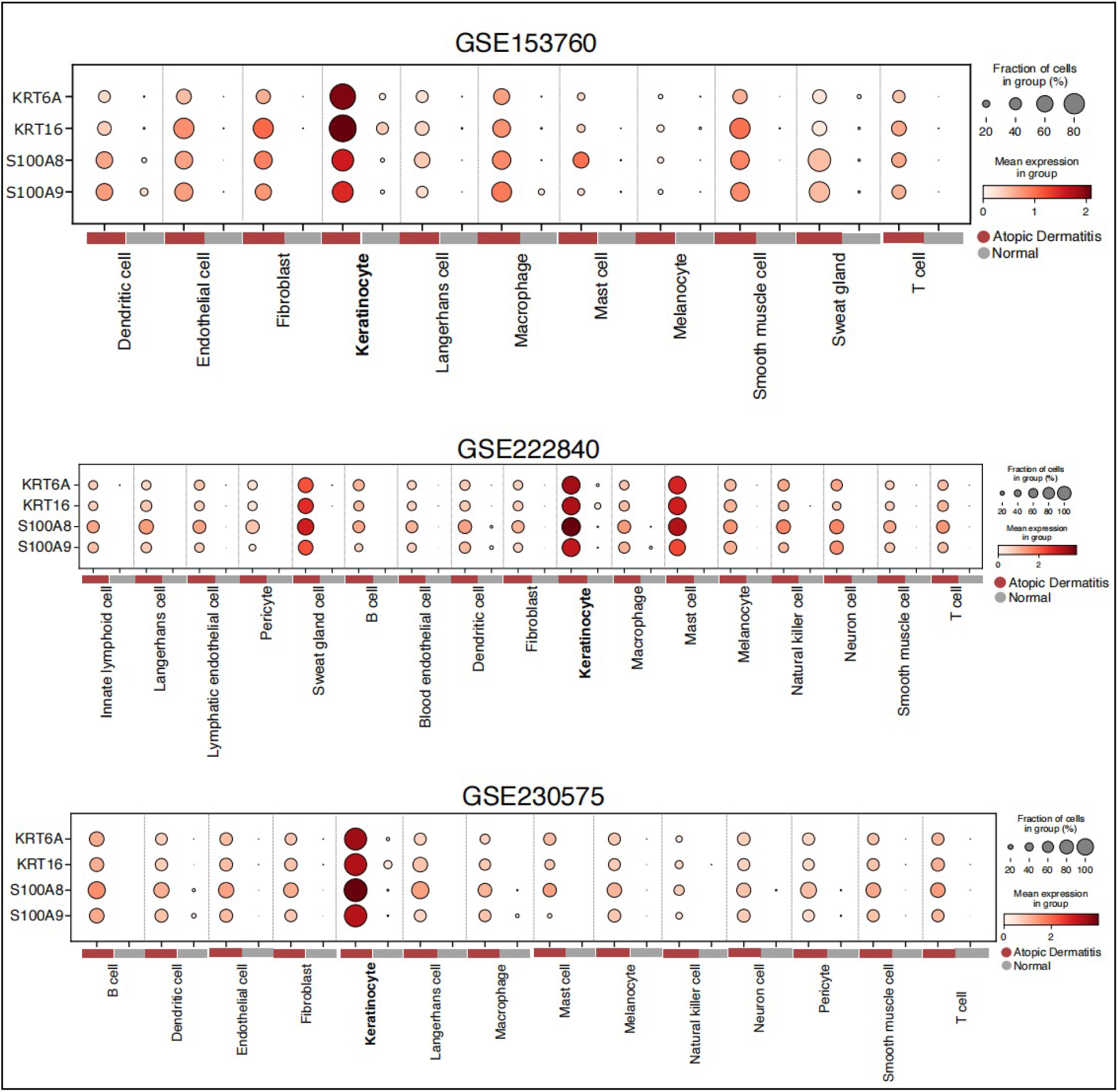
Comparison of KRT6A, KRT16, S100A8, and S100A9 expression between AD and healthy samples. The comparisons were run on 3 individual studies and on each cell type separately. All cell types showed consistently higher expression of all 4 genes in AD compared to healthy samples, with the highest significance in keratinocytes.

**Supplementary Figure 3.**
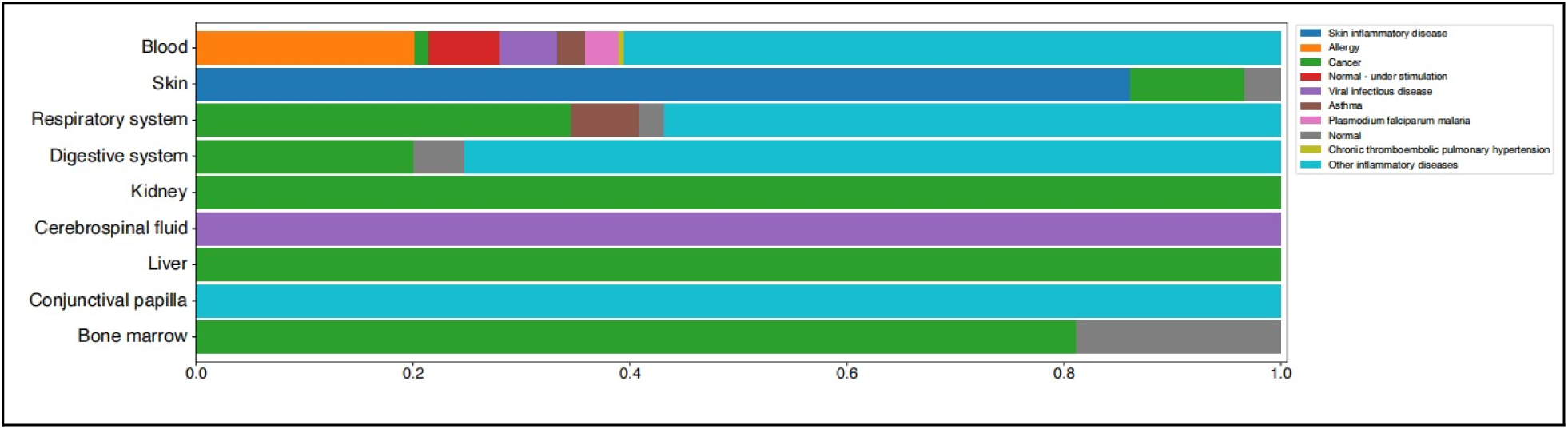
Proportions of conditions within each tissue.

**Supplementary Figure 4.**
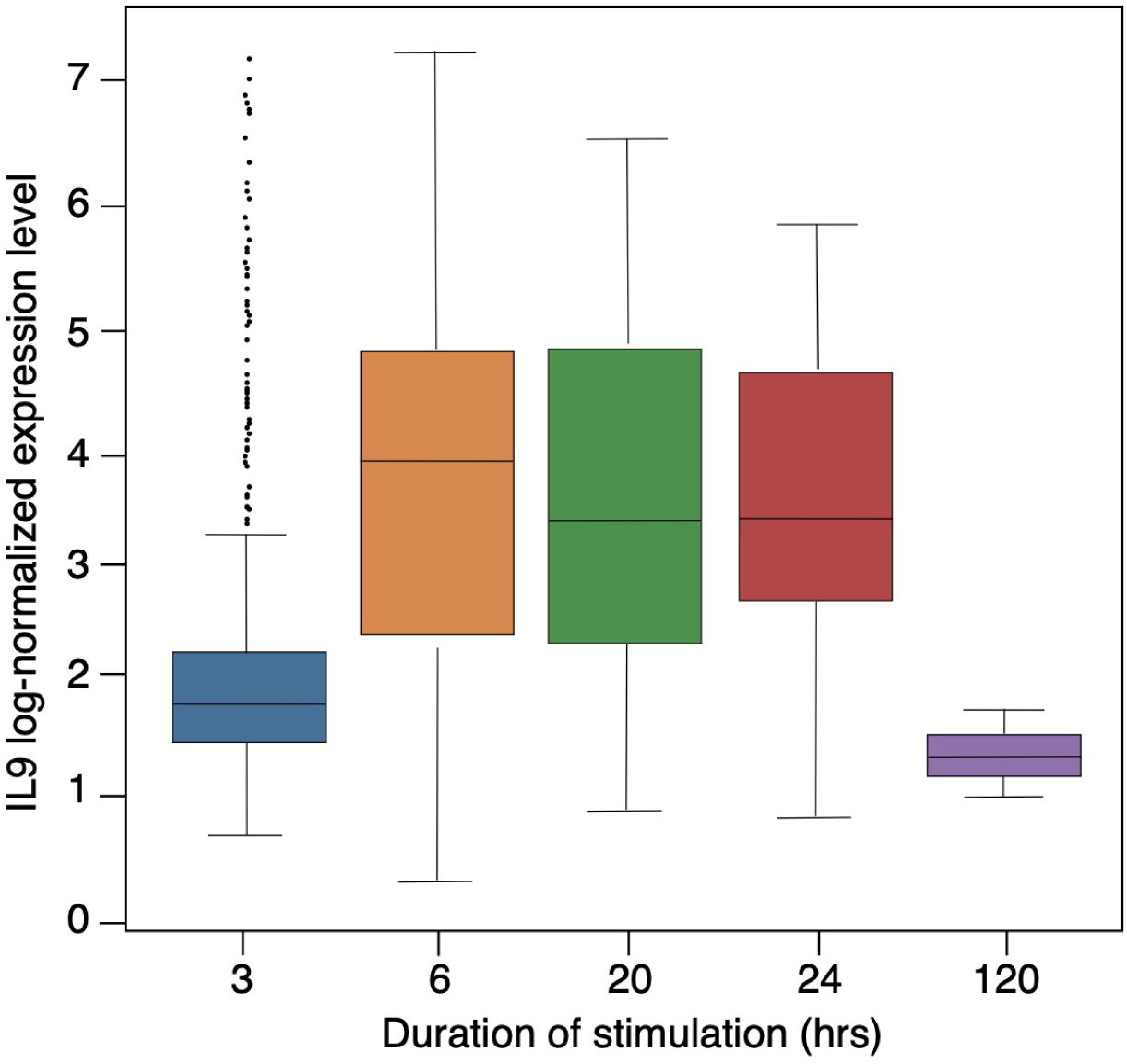
Box plot of IL9 normalized expression across different allergen contact/stimulation durations. Across different allergic conditions in the current CD4+Th2 atlas. 3hrs: Food allergy (blue; GSE136028_HUMAN); 6hrs: EE and A/ARD (orange; GSE175930, GSE146170); 20hrs: Food allergy (green; GSE158667); 24hrs: A/ARD (red, GSE193816, GSE164059); 120hrs: allergic rhinitis (purple; GSE200107).

Supplementary Table 1:

**Datasets included in the CD4+ Th2 Atlas Th2 datasets**

